# Incidence and susceptibility patterns of urine bacterial flora in young females

**DOI:** 10.1101/2020.03.18.996793

**Authors:** Ashwag Shami, Samiah Al-Mijalli, Ali Somily, Fawziah M. Albarakaty, Samah Awad AbduRahim

**Author notes:** **Corresponding author:** Samah Awad AbduRahim,.

## Abstract

**Background:** It has been established that the urinary tract is not sterile, however, research related to the study of urine bacteria are limited. Our work aims to study the frequency and patterns of resistance of normal urinary aerobic bacterial flora.

**Methods:** Clean catch midstream urine specimens were collected from 120 young healthy females, and then cultured. Identification of bacteria and antimicrobial susceptibility were done by means of Biomérieux VITEK® 2 automated systems. Subjects who have undergone antimicrobial treatment in one month weren’t included.

**Results:** The incidence of positive bacterial cultures was 54.2%, of which 21.5% were ploymicrobial. 107 bacterial isolates that encompass 12 genera and 27 species that were predominated by Gram-positive bacteria (72%) were cultivated. Staphylococcaceae (46.1%) and Enterobacteriaceae (17.8%) were the most frequently isolates among Gram positive and Gram negative bacteria respectively, from them 36 species have been identified as b lactamase producers. The top four frequently isolated bacteria are Micrococcus species (16%), *Staphylococcus haemolyticus* (13.2%), *Staphylococcus aureus* (10%), and *Klebsiella pneumoniae* (10%). Twenty two bacterial species were subjected to antimicrobial susceptibility testing by using broad and narrow spectrum antibiotics and antimicrobials. Ampicillin showed the lowest susceptibility rate against Gram positives, followed by erythromycin and azithromycin. The lesser antimicrobial susceptibility potential among Gram negative bacteria was exhibited against ampicillin, followed by piperacillin and cefotaxime.

**Conclusion:** Our findings emphasize the importance of highlighting urine bacterial flora in research especially those related to susceptibility patterns, employing more advanced culture methods as multiple drug resistant bacteria were isolated.

## Introduction

The recent researches have emphasized the residence of bacteria in the urine of women; the fact decided that the use of antibiotics based on a positive culture should be accompanied by clinical symptoms [1, 2]. For the time being, culture methods were regarded by some microbiologists ineffective in reflecting bacterial diversity in the human body as do some technological methods that were developed after the tremendous progress in the genomics, which have turned out unprecedented revolution in deeply recognition of body flora [3]. However, despite their limitations in identifying many types of bacteria, as well as they are accused of being the cause of the ancient prevailing concept which states that the urinary tract is not occupied naturally by bacteria, notwithstanding, the culture methods remain with all strength a number that cannot be exceeded due to their importance in identifying patterns of bacterial susceptibility to the antimicrobial agents which represents the initial phase in setting up suggestions for powerful treatment for infectious diseases [4, 5]. Therefore, unsurprisingly, Robert Koch considered them are the fundamental of infectious disease research [6]. Under usual conditions; normal flora did not represent a danger to the host, but recent studies reported that these bacteria are possessed or can extrinsically acquire some genes responsible for resistance to some antibiotics have sounded the alarm bell for this emergency and drew attention to the necessity of conducting susceptibility testing for them [7, 8, 9]. After tracing and extrapolation of the research that interested in bacterial flora, it became clear to us that information is very scarce about susceptibility patterns of these microorganisms, especially those that settle urinary tracts. Therefore, this study was conducted to investigate the urinary flora in young healthy female and to assess the antibiotic susceptibility of recovered bacteria.

## Methodology

### Study population

Participants enrolled in the study were unmarried Saudi female between the ages of 21-23 years recruited at Princess Nourah bint Abdulrahman University and King Khalid hospital (n=120). The female haven’t any history of overactive bladder syndrome, or urinary incontinence, urgency, or frequency.

### Study area and specimen collection

The study was conducted in Princess Nourah bint Abdulrahman University and King Saud University Research Centre in Riyadh. After collection, the midstream clean catch urine specimen was aliquoted into sterile 15 mL-conical tubes and stored at −20°C.

### Ethical approval

All procedures and techniques in this study were in accordance with national regulations that govern the protection of human subjects. The protocols were approved by Princess Nourah bint Abdulrahman Institutional Review Board (under IRB number: 17-0060). Urine specimen was collected after obtaining a signed informed consent from the participants. All participants recruited were older than 16 years old.

### Specimen culture and bacterial identification

Urine culture was performed by inoculating 200 μl of urine onto the entire plate of blood agar plate. The plates were incubated at 37°C under aerobic condition overnight. Identification of bacteria and antibiotic susceptibility were carried out adopting Biomérieux VITEK® 2.

### Data analysis

The data were analysed using SPSS (version 21), and Microsoft Excel 2010 soft wares. Descriptive statistics were performed for numbers of susceptible and resistant bacterial species; the results were expressed as mean ± standard deviation. Kruskal-Wallis test was used to determine the statistical differences between the means resistance of antibiotics. Mann-Whitney test was done for each pair of antibiotics and bacteria to decide which one causes the difference. The mean difference is significant at the 0.05 level.

## Results

There were 120 female participants recruited at Princess Nourah bint Abdulrahman University and King Khalid hospital, those in the age group between 20-25 years old, none of them showed signs or symptoms of urinary tract infection. It was observed that 45.8% of the investigated midstream urine specimens were sterile. Positive cultures were obtained from 65 specimens (54.2%), of which 14 were ploymicrobial. 107 bacterial isolates that encompass 12 genera and 27 species that were predominated by Gram-positive bacteria which represent (72%) were cultivated. Staphylococcaceae (46.1%) and Enterobacteriaceae (17.8%) were the most frequently isolates among Gram positive and Gram negative bacteria respectively. The frequency, percentage and numbers of the isolated urinary flora are presented in table (1) and table (2). Of 10 isolates, two *S. aureus* strains were identified as methicillin-resistance *S. aureus* (MRSA); however, they remained susceptible to vancomycin. It is worth noting that 36 b-lactamase producing bacteria have been identified among Gram positives. Twenty two bacterial species were subjected to antimicrobial susceptibility testing by using broad and narrow spectrum antibiotics and antimicrobials. We described the susceptibility patterns as resistant when 100 % of tested bacteria were resistant, relatively resistant when only part of them when not all bacteria are resistant, but rather there are sensitive and resistant species, and susceptible when 100 % of tested bacteria were susceptible. The mean numbers of susceptible, relatively resistant and resistant species among Gram negatives were 1.19 ± 0.81, 1.19 ± 1.25, and 3.86 ± 2.17 respectively. On the other hand, the mean numbers of susceptible, relatively resistant and resistant species among Gram positives were 2.63 ± 1.92, 3.68 ± 2.26, and 6.42 ± 3.49 respectively.

**Table 1:** The profile, percentage and numbers of the isolated Gram-positive bacteria.

| Genus | Species | No. of the isolates | (%) |
| --- | --- | --- | --- |
| **Enterococcus** | *Enterococcus faecalis* | 3 | 2.8 |
|  | *Enterococcus faecium* | 3 | 2.8 |
| **Kocuria** | *Kocuria kristinae* | 1 | 0.9 |
| **Micrococcus** | *Micrococcus* | 17 | 16 |
| **Rothia** | *Rothia dentocariosa* | 3 | 2.8 |
|  | *Rothia mucilaginosa* | 1 | 0.9 |
| **Staphylococcus** | *Staphylococcus aureus* | 10 | 9.4 |
|  | *Staphylococcus auricularis* | 2 | 1.9 |
|  | *Staphylococcus cohnii* | 1 | 0.9 |
|  | *Staphylococcus epidermidis* | 6 | 5.7 |
|  | *Staphylococcus haemolyticus* | 14 | 13.2 |
|  | *Staphylococcus hominis* | 2 | 1.9 |
|  | *Staphylococcus hyicus* | 1 | 0.9 |
|  | *Staphylococcus intermedius* | 2 | 1.9 |
|  | *Staphylococcus lugdunensis* | 3 | 2.8 |
|  | *Staphylococcus sciuri* | 6 | 5.7 |
|  | *Staphylococcus simulans* | 1 | 0.9 |
|  | *Staphylococcus warneri* | 1 | 0.9 |

**Table 2:** The profile, percentage, and numbers of the isolated Gram-negative bacteria.

| Genus | Species | No. of the isolates | (%) |
| --- | --- | --- | --- |
| **Acinetobacter** | *Acinetobacter baumannii* | 2 | 1.9 |
| **Cedecea** | *Cedecea lapagei* | 1 | 0.9 |
| **Citrobacter** | *Citrobacter murlinae* | 1 | 0.9 |
| **Escherichia** | *Escherichia coli* | 7 | 6.6 |
| **Klebsiella** | *Klebsiella pneumoniae* | 10 | 9.4 |
| **Pseudomonas** | *Pseudomonas aeruginosa* | 5 | 4.7 |
|  | *Pseudomonas fluorescens* | 1 | 0.9 |
|  | *Pseudomonas stutzeri* | 1 | 0.9 |
| **Burkholderia** | *Burkholderia cepacia* | 1 | 0.9 |

Figure (1) and (2) describe the numbers and patterns of susceptibility of Gram-positive and Gram-negative bacteria to antibiotics used for each species. Table (3) shows the patterns of susceptibility towards seven antibiotics simultaneously tested against both types of bacteria.

**Figure (1):**
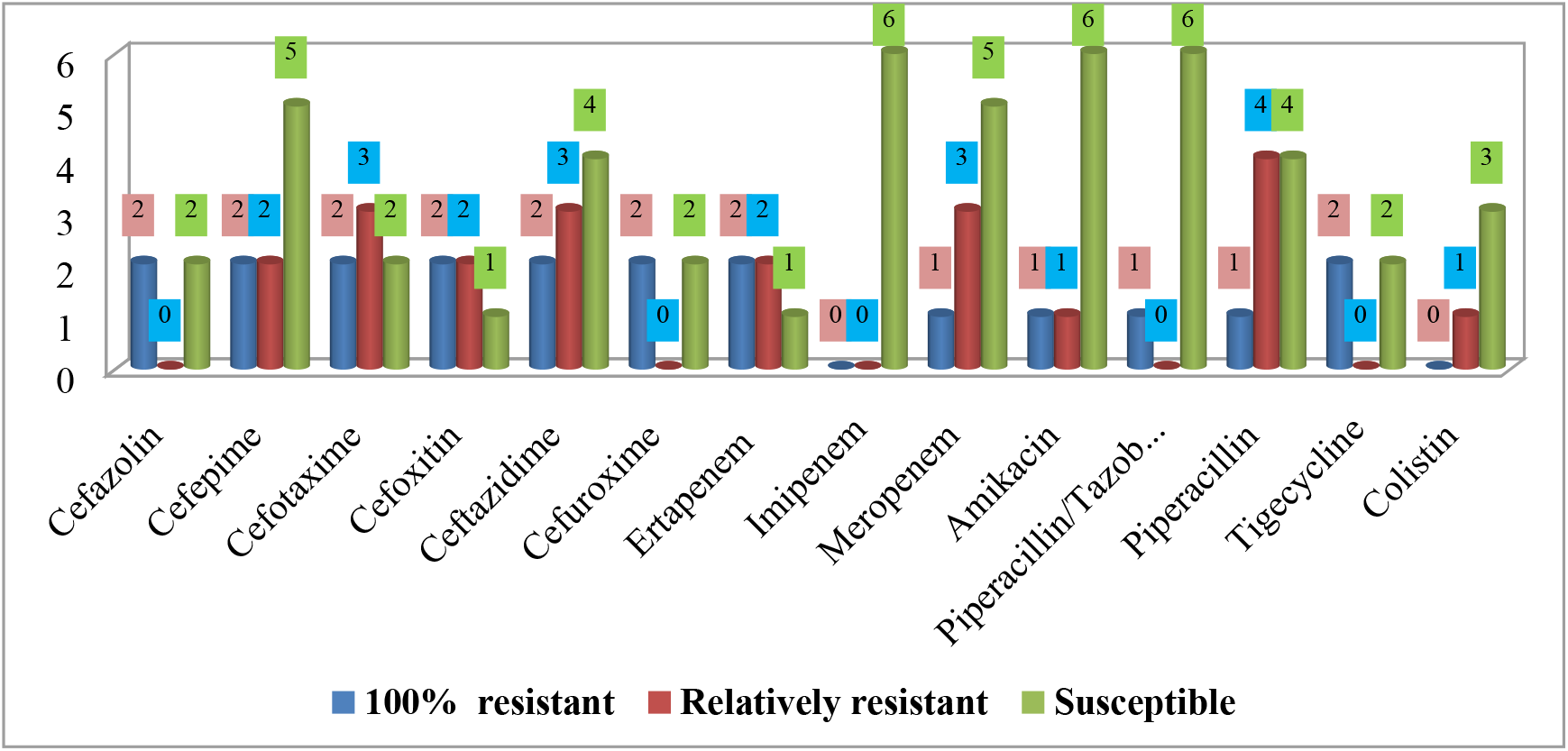
Numbers and susceptibility patterns of Gram-negative urinary flora

**Figure (2):**
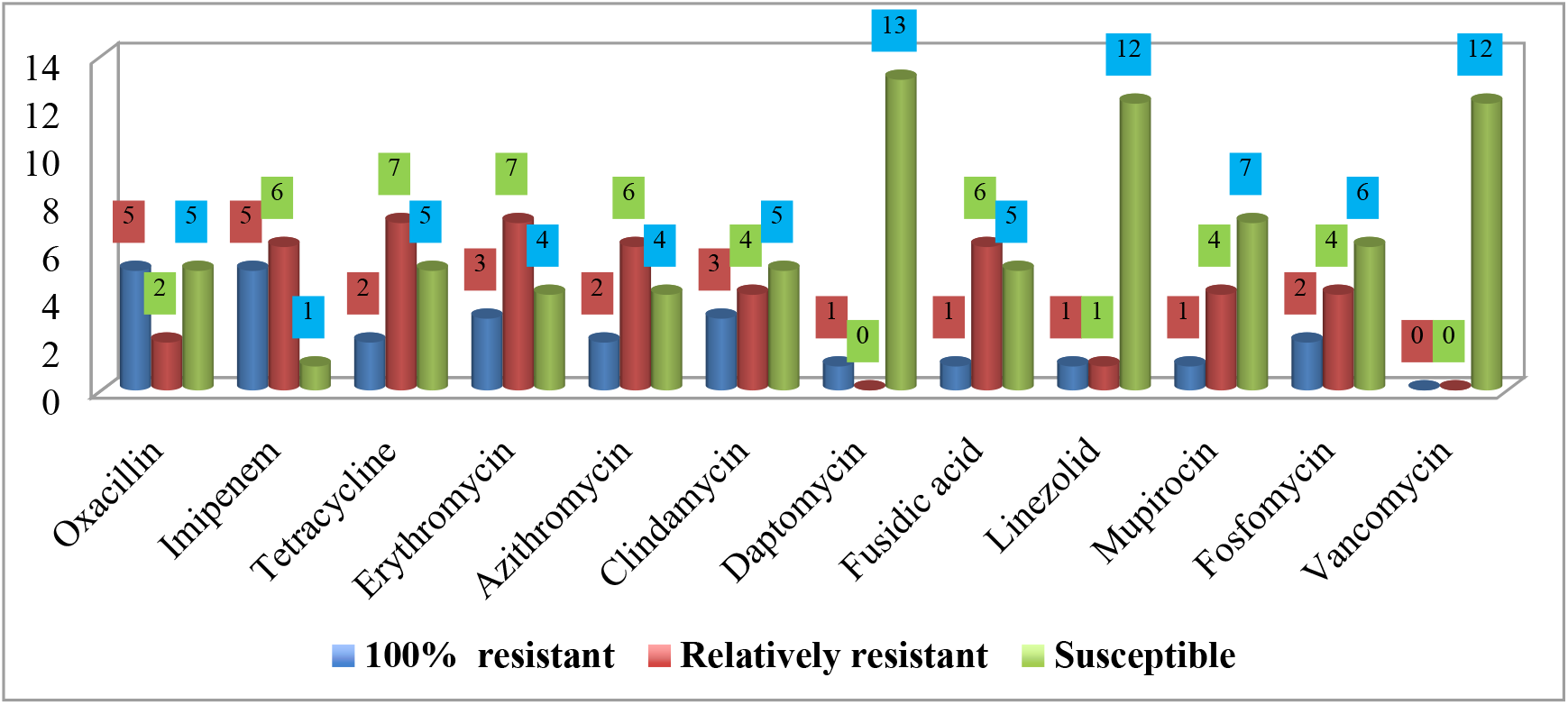
Numbers and susceptibility patterns of Gram-positive urinary flora

**Table 3:** Susceptibility patterns to the antibiotics tested against both bacterial types.

|  | 100% resistant |  | Relatively resistant |  | Susceptible |  | Total |  |
| --- | --- | --- | --- | --- | --- | --- | --- | --- |
| Antimicrobial agent | Gram negative | Gram positive | Gram negative | Gram positive | Gram negative | Gram positive | Gram negative | Gram positive |
| **Amoxicillin/Clavulanic acid** | 2 | 5 | 0 | 6 | 2 | 1 | 4 | 12 |
| **Ampicillin** | 0 | 8 | 1 | 3 | 3 | 3 | 4 | 14 |
| **Gentamicin** | 1 | 2 | 1 | 4 | 6 | 6 | 8 | 12 |
| **Levofloxacin** | 0 | 2 | 0 | 2 | 8 | 10 | 8 | 14 |
| **Ciprofloxacin** | 0 | 3 | 0 | 3 | 8 | 8 | 8 | 14 |
| **Moxifloxacin** | 1 | 2 | 1 | 1 | 2 | 9 | 4 | 12 |
| **Sulfamethoxazole/Trimethoprim** | 1 | 2 | 1 | 4 | 3 | 6 | 5 | 12 |

Ampicillin showed the lowest susceptibility rate against Gram positives, followed by erythromycin and azithromycin. High susceptibility rates were reported to vancomycin (0%), daptomycin (1.3%), and linezolid (2.6%). The susceptibility profile of Gram-positive bacteria for 19 antibiotic agents was summarized in Table (4).

**Table 4:**
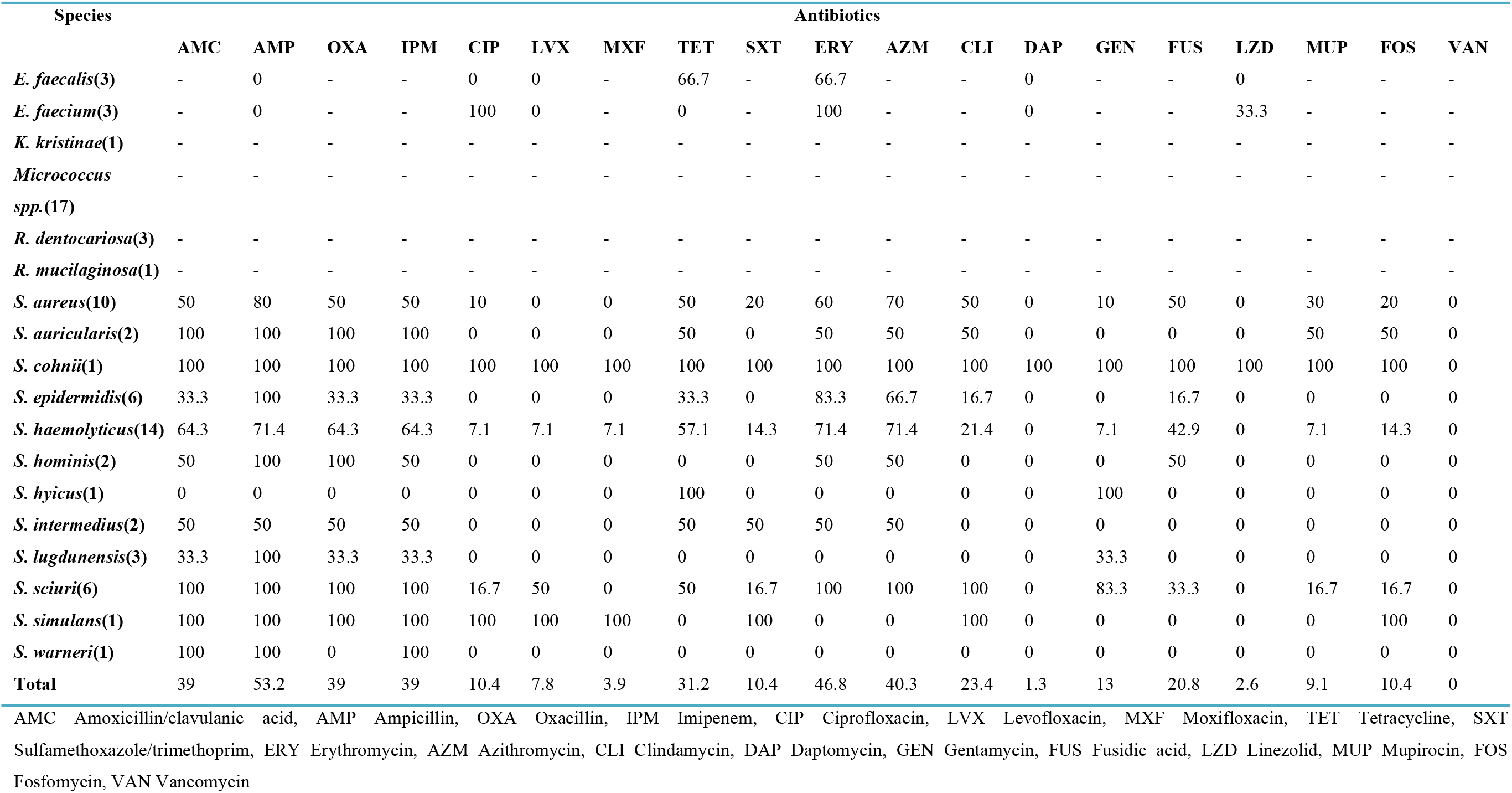
The susceptibility profile of Gram-positive bacterial flora.

The susceptibility profile of Gram-negatives for 22 antibiotic agents was summarized in Table (5). The lesser antimicrobial potential was exhibited against ampicillin, followed by piperacillin and cefotaxime. Ciprofloxacin and levofloxacin remained active, with the resistance rate of 0%.

**Table 5:** Table 4: The susceptibility profile of Gram-negative bacterial flora.

| Species | Antibiotics |  |  |  |  |  |  |  |  |  |  |  |  |  |  |  |  |  |  |  |  |  |
| --- | --- | --- | --- | --- | --- | --- | --- | --- | --- | --- | --- | --- | --- | --- | --- | --- | --- | --- | --- | --- | --- | --- |
|  | AMC | AMP | CFZ | CFP | CTX | FOX | CAZ | CXM | ETP | IPM | MEM | AMK | GEN | LFX | CIP | MXF | TZP | PIP | TET | TGC | SXT | CST |
| *A. baumannii*(2) | - | - | - | 0 | 50 | - | 0 | - | - | - | 0 | 0 | 0 | 0 | 0 | - | - | 50 | 0 | - | 0 | - |
| *C. lapagei*(1) | 100 | 100 | 100 | 100 | 100 | 100 | 100 | 100 | 100 | 0 | 0 | 100 | 100 | 0 | 0 | 0 | 0 | 0 | 100 | 100 | 100 | - |
| *C. murlinae*(1) | 100 | 100 | 100 | 100 | 100 | 100 | 100 | 100 | 100 | - | 0 | 0 | 0 | 0 | 0 | 100 | 100 | 100 | 100 | 100 | 0 | - |
| *E. coli*(7) | 0 | 14.3 | 0 | 0 | 0 | 0 | 0 | 0 | 0 | 0 | 0 | 0 | 0 | 0 | 0 | 14.3 | 0 | 14.3 | 14.3 | 0 | 0 | - |
| *K. pneumoniae*(10) | 0 | 100 | 0 | 0 | 0 | 10 | 10 | 0 | 10 | 0 | 10 | 0 | 0 | 0 | 0 | 0 | 0 | 40 | 20 | 0 | 10 | 0 |
| *P. aeruginosa*(5) | - | - | - | 20 | 60 | - | 20 | - | - | 0 | 20 | 20 | 20 | 0 | 0 | - | 0 | 0 | - | - | - | 20 |
| *P. fluorescens*(1) | - | - | - | 0 | - | - | 0 | - | - | 0 | 100 | 0 | 0 | 0 | 0 | - | 0 | 0 | - | - | - | 0 |
| *P. stutzeri*(1) | - | - | - | 0 | - | - | 0 | - | - | 0 | 0 | 0 | 0 | 0 | 0 | - | 0 | 0 | - | - | - | 0 |
| *B. cepacia*(1) | - | - | - | - | - | - | - | - | - | - | - | - | - | - | - | - | - | - | - | - | - | - |
| **Total** | 6.9% | 44.8% | 6.9% | 10.3 | 20.7 | 10.3 | 13.8 | 6.9% | 10.3 | 0.0% | 10.3 | 6.9% | 6.9% | 0.0% | 0.0% | 6.9% | 3.4% | 24.1 | 17.2 | 6.9% | 6.9% | 3.4% |
|  |  |  |  | % | % | % | % |  | % |  | % |  |  |  |  |  |  | % | % |  |  |  |

AMC Amoxicillin/Clavulanic acid, AMP Ampicillin, CFZ Cefazolin, CFP Cefepime,CTX Cefotaxime, FOX Cefoxitin,CAZ Ceftazidime, CXM Cefuroxime, ETP Ertapenem, IPM Imipenem, MEM Meropenem, AMK Amikacin, GEN Gentamicin, LFX Levofloxacin, CIP Ciprofloxacin, MXF Moxifloxacin, TZP Piperacillin/Tazobactam, PIP Piperacillin, TET Tetracycline, TGC Tigecycline, SXT Sulfamethoxazole/Trimethoprim, CST Colistin

## Discussion

The natural bacteria in the urinary tract did not receive extensive research, as with the skin and gut bacteria. In addition to the fact that they were not included in the human microbiome project [4], they did not obtain enough research in their antimicrobial susceptibility patterns, and this may be because urinary system was formerly deemed to be a sterile niche [10, 11]. However, recent studies have shown that it is naturally colonized by a variety of bacterial species [12], and it has become certain that their lopsidedness -somehow- have a role in the system’s physiology and susceptibility to infection as they represent a biological defence line against “pathogen induced inflammation” [13, 14]. Numerous studies have confirmed the difference in bacterial communities between healthy and women with urgency urinary incontinence (UUI) [15, 16]. Thomas et al and Mueller et al correlated between the response to the treatment in patients with UUI and dysbiosis of the urinary flora [17, 18].

In the present study, antibiotics have not undergone susceptibility tests against Micrococci, Rothiae*, Kocuria kristinae*, and *Burkholderia* cepacia. This is because automated identification systems - although they allow accurate identification-unfortunately, do not perform susceptibility tests for some bacterial species, the fact that remains one of the most important deficiencies of these systems [19, 20]. There are not adequate recent studies to indicate the pattern of Micrococcal response to antibiotics, but it is convenient here to refer to the Dürst etal findings which showed the response of *Micrococcus luteus* to penicillin [21]. Scarce information is available on the pathogenesis of Micrococcus species as well, but in general they seldom produced beta-lactamases [22], and it is difficult to attribute the aetiology of the infection to this genus if it is isolated from a clinical specimen as it rarely affects people with healthy immune system, and the high virulence remains limited to people with low immunity, such as AIDS patients [23, 24]. Nevertheless, they have given rise to some infections latterly such as pneumonia, meningitis, septic arthritis, peritonitis and endophthalmitis [25–29]. We isolated 5 genera out of the six earnest multiple drug resistant (MDR) bacteria which abbreviated by “ESKAPE” referring to their acronyms. ESKAPE encompass *Enterococcus faecium*, *Staphylococcus aureus*, *Klebsiella pneumoniae*, *Acinetobacter baumannii*, *Pseudomonas aeruginosa* and Enterobacter spp [30].

14.3 % of *E.coli* isolates were resistant to ampicillin, piperacillin, tetracyclin and moxifloxacin, however they were inhibited by penicillin agents combined with b lactamase inhibitors (amoxicillin/cavulanicacid - amoxiclav- and piperacillin/ tazobactam (TZP)). None of the isolates was resistant to imipenem, tigecycline and colistin, similar to findings of Rasheed et al [31], who reported the resistance to ampicillin and tetracycline. However he documented resistance to cefotaxime (5.3%), gentamycin and amoxicillin/cavulanic acid (4.6 %) in contention of our results where no resistance was encountered the above mentioned antibiotics. Out of 11 antibiotics, A*. baumannii* isolates were susceptible to 9 antibiotics and only 50% of isolates were resistant to cefotaxime and piperacillin, and these outcomes are good as the bacterium has emerged as an opportunistic MDR pathogen associated with nosocomial infections [32], but to best of our knowledge no literature was available in correspondence to the susceptibility of urinary *A. baumannii* flora. All *Klebbsiella pneumoniae* strains were resistant to ampicillin and susceptible to amoxiclav. This might suggest that the mechanism of resistance may refer to TEM-1; an enzyme mediates ampicillin resistance and can be inhibited by b-lactamase inhibitors [33], the same reason that can explicate the susceptibility of *Klebsiella pneumoniae* isolates to TZP. Osman et al contradicted our results when they reported colistin and ampicillin resistance among *K. pneumniae* strains isolated from milk, nevertheless, similarly to the recent outcomes, no resistance was demonstrated to gentamycin, cefatoxime, and slphmethoxazole/trimethoprim (SXT) [34]. One strain exhibited resistance to ertapenem and meropenem, this is alarming since genes encoding the *Klebbsiella pneumonia* carbapenemase (KPC) and New Delhi Metallo b lactamase (NDM) are harbored in plasmids, thus can be transmitted to other species [35]. The susceptibility patterns of *Pseudomonas fluorescens* and *Pseudomonas stutzeri* were found to be similar in exception of that the former was resistant to meropenem. Kittinger et al reinforced our susceptibility results about meropenem against *P. fluorescens*, but contrary, he reported resistance to ceftazidime and ciprofloxacin [36]. *P. stutzeri* was susceptible to the 19 antibiotics tested identically to its susceptibility results when isolated from a patient with peritonitis [37]. Park et al attributed the high susceptibility of this bacterium to its rare clinical incidence and thus less exposure to antimicrobials [37]. Out of 21, *Cedecea lapagei* was resistant to 14 antibiotics including amikacin, gentamycin, ceftazidime, cefepime, and sulfamethoxazole/trimethoprim, an outcome contradicted with Biswal etal, because they reported the bacterium susceptibility to above mentioned antibiotics [38]. This might give a prime example of bacterial adaptation to the antibiotics through acquisition of resistance determinants [39, 33]. However, our findings are in agreement with Biswal etal as they demonstrated the bacterium susceptibility to meropenem and ciprofloxacin along with its resistance to tetracycline and tigecycline [38]. *Citrobacter murliniae* was identified as an operational taxonomic unite in female urine in 2017 [40], but no literature was available – to our knowledge-about its susceptibility profile. It expressed multidrug resistant phenotype similarly to *C. lapagei* (Table 5). Highlighting Gram positives, the two Enterococci were susceptible to ampicillin, levofloxacin and daptomycin. Closely similar percentage to penicillin (96%) was reported by Rudy et al in their screening of *Enterococcus faecalis* isolated from urine, but they observed a higher resistant rate to ciprofloxacin (43%) [41].They found that 14% of *Enterococcus faecium* strains were resistant to ciprofloxacin, 32 % to ampicillin, and 19 % to tetracycline. Our study remained consistent with previous studies that showed that urinary tract is predominantly inhabited by coagulase negative Staphylococci (CONS) [42, 43]. *S. haemolyticus* was documented as a part of female urinary normal flora [44], congruously; Pindar et al reported the organism’s susceptibility to vancomycin [45].

*Staphylococcus simulans* was found to be resistant to the all b-lactams and quinolones. It was isolated by Shields et al as a skin associated pathogens [46], however, sparsely data is available about the antimicrobial susceptibility of this emerging pathogen. We didn’t recover *S. saprophyticus* although it is associated with community acquired urinary tract infections, secondly after *Escherichia coli* [43, 47]. To best of our knowledge, it was not isolated from the mid-stream urine except in low percentages ranged between 2%-4% in sexually active females and pregnant women [48–51], where it was encountered in low bacteriuria [52]. The difference in the prevalence percentages can also be attributed to the fact that the prevalence of *S. saprophyticus* changes seasonally and increases significantly at the end of summer [53]. We isolated three coagulase positive Staphylococci, *S.aureus, S. intermedius, and S. hyicus* (which was found to be MDR), but it is surprising that the second and third are related to dogs and pigs respectively [54, 55], and there is no evidence, to the best of our knowledge, that it is part of the natural components of the human’s urinary system. *S. aureus* exhibited significantly lower susceptibility (P≤0.05) to tetracycline, erythromycin, azithromycin, clindamycin, mupirocin, and fusidic acid than CONS. However, the latter demonstrated higher resistant rate to the remaining antibiotics consistently with Tao et al reported outcomes [56]. Comparing to CONS with resistance rate average, 20.3%, *S. aureus* showed lower resistance rate with 0.0–10 % to quinolone. Likewise, vancomycin and linezolid don’t encounter resistance among *S. aureus* isolates. This also emphasized the compatibility with Tao et al findings.

## Conclusion

Our study concludes that the number of isolated aerobic bacterial flora in the clean catch mid-stream urine was significantly higher in gram positives. It was a good indication that Gram-positive bacteria, including MRSA and other b-lactamase producers, did not show any resistance at all to vancomycin. Imipenem, levofloxacin, and ciprofloxacin were completely active against Gram-negative bacteria. On the other hand, it is of concern that there is resistance to some antibiotics proved to be mediated by mobile genetic elements that can be transmitted between bacterial species. Certainly, these results are not final; rather, we recommend more research in this field by employing modern methods of culture such as matrix assisted laser desorption ionization-time of flight mass spectrometry (MALDI-TOF MS) to isolate broader spectra of bacteria, ensuring a deeper study of susceptibility and resistance patterns of urine bacterial flora.

## Acknowledgement

We would like to thank all the staffs of Unit of Microbiology, King Saud Hospital University, and Department of Microbiology, Princess Nourah Bint Abdulrahman University for their hard work.

## Conflict of interest

No conflict of interest to declare

## Funding

This research was funded by the Deanship of Scientific Research at Princess Nourah bint Abdulrahman University through the Fast-track Research Funding program.

